# Carryover effects of tire wear particle leachate threats reproduction across multiple generations

**DOI:** 10.1101/2022.10.27.513999

**Authors:** Yanchao Chai, Haiqing Wang, Mengru Lv, Jiaxin Yang

## Abstract

Toxic additives leached from tire wear particle (TWP) have been linked to some collective death events of fish, also impose impacts on zooplankton as secondary consumer in aquatic food web. In addition to direct impacts of TWP leachate at the current generation, potential delayed carryover from past exposure across multi-generational lineage may augment impacts on individual reproduction, then population maintenance. We investigated the carryover effects from persistent exposure and past TWP leachate exposure along generation passaging on the individual reproduction of rotifer, *Brachionus calyciflorus*, a typical zooplankton. For rotifer treated with TWP leachate across continuous 7 generations, their offspring were divided into parental exposure or no exposure in each generation. And rotifer transferred into no exposure were maintained for 3 generations to eliminate indirect exposure through their parental germ. The similar response of reproduction, via carryover effects from parental exposure, also emerged in offspring without exposure. Persistent exposure across multiple generations additively impaired individual reproduction performance by transferring from its hermetic effects in original generation, even caused final population collapse. TWP leachate could impose cascading toxicity on population persistence of zooplankton via carryover and cumulative effects on reproduction for long term, which must be considered in risk assessment and management policy to alleviate the effects of TWP.

## 1. Introduction

Sporadic collective deaths of salmon in urban stream have been linked to the extractable additive from tire wear particle (TWP) in road runoff (Tian et al., 2021), which should be a wake-up call about the effects of TWP on aquatic ecosystem function and stability. Substantial increasing of global tire utilization, as well as complex toxic additives in them and non-point source pollution without effective control, emphasize the urgency to understand the magnitude of impacts of TWP leachate on aquatic biome (Halsband et al., 2020; Knight et al., 2020; Kole et al., 2017; Wagner et al., 2018). Studies of these impacts have primarily focused on the effects of TWP leachate at just current generation exposure. However, TWP leachate may substantially affect the production of offspring long after exposure, or its consequences were magnified through persistent exposure across multiple generations.

Those carryover effects across generations, in which past experience impacts individual performance of current generation, are observed in some taxa exposed to other stressors (Stuligross and Williams, 2021). That is also referred to transgenerational and multigenerational effects (Jeremias et al., 2018; Patel et al., 2019; Trijau et al., 2018). For toxic substances, they can accumulate *in vivo* and aggregate damages after continuous exposure of multiple generations (Guo et al., 2012). Even if the exposure is eliminated, the effects can be imprinted in offspring due to toxicity residual or epigenetic modification along germ linage (Ellis et al., 2020; Jeremias et al., 2018; Li et al., 2015; Trijau et al., 2018; Vandegehuchte et al., 2010). Those delayed impacts can be ignored in current exposure on single generation, which underestimates effects of stressors.

For some aquatic invertebrates, the flourishing of population depends on the early pioneers, which then goes through quick generation passaging. Carryover effects from the exposure on the original generation may disturb population dynamics. Meanwhile, they lack strong swimming mobility to escape local stressors, and are also sensitive to exogenous stressors. Carryover effects from some contaminants have been recorded, but the effects of TWP leachate have not been explored.

Some studies have shown that the maternal exposure of some heavy metals and organic compounds, as main components of TWP leachate, could have carryover effects on offspring (Li et al., 2015; Paumen et al., 2008). That includes that reduced resistance to stressors, lifespan and production, which are vital for population maintenance. The invasion of TWP leachate is similar to impulse-style discharge pattern due to accidental road storm runoff. It means that the exposure of TWP leachate could just happens temporarily at limited life stage or generations. Thus, it is necessary to understand if this temporary exposure influence reproduction via carryover effects in long term.

Understanding these delayed effects of TWP leachate is crucial for population maintaining of zooplankton, as a vital intermediate link in food web, which guarantee stability of production and services in aquatic ecosystem. The negative effects of TWP leachate have been well documented on some zooplankton, such as Cladocera and Copepods (Wik and Dave, 2006). Some sublethal effects are observed, including reduced reproduction and lifespan, but carryover effects have not been verified.

We investigated the carryover effects of TWP leachate exposure on the individual reproduction of the rotifer, *Brachionus calyciflorus*. The offspring from each generation experienced TWP leachate were persistently exposed (or not) across 7 generations (P-F6). And the individuals with no more exposure were breed to 3^th^ generations to avoid the exposure through germ line. The offspring number born from individual was used to assess reproduction performance. We obtained TWP from factory of scrap tire recycle, with broad representation for wide variety of tires. Using a multigenerational experiment with persistent exposure and recovery line (no more exposure; Fig. 1), it can be explored that 1) whether the TWP exposure on past generations can carry over to additively influence reproduction of current generation, 2) whether the toxicity is alleviated after multi-generational exposure when evident damage is triggered in original generation, 3) whether reproductive damages can carry over to influence rotifers without exposure. Intergenerational accumulation, adjustment or heritage of toxicity of TWP leachate on reproduction of *B. calyciflorus* will be verified. The utility of multigenerational toxicity tests in this study may predict more accurately effect threshold of TWP leachate in long term.

**Fig. 1.**
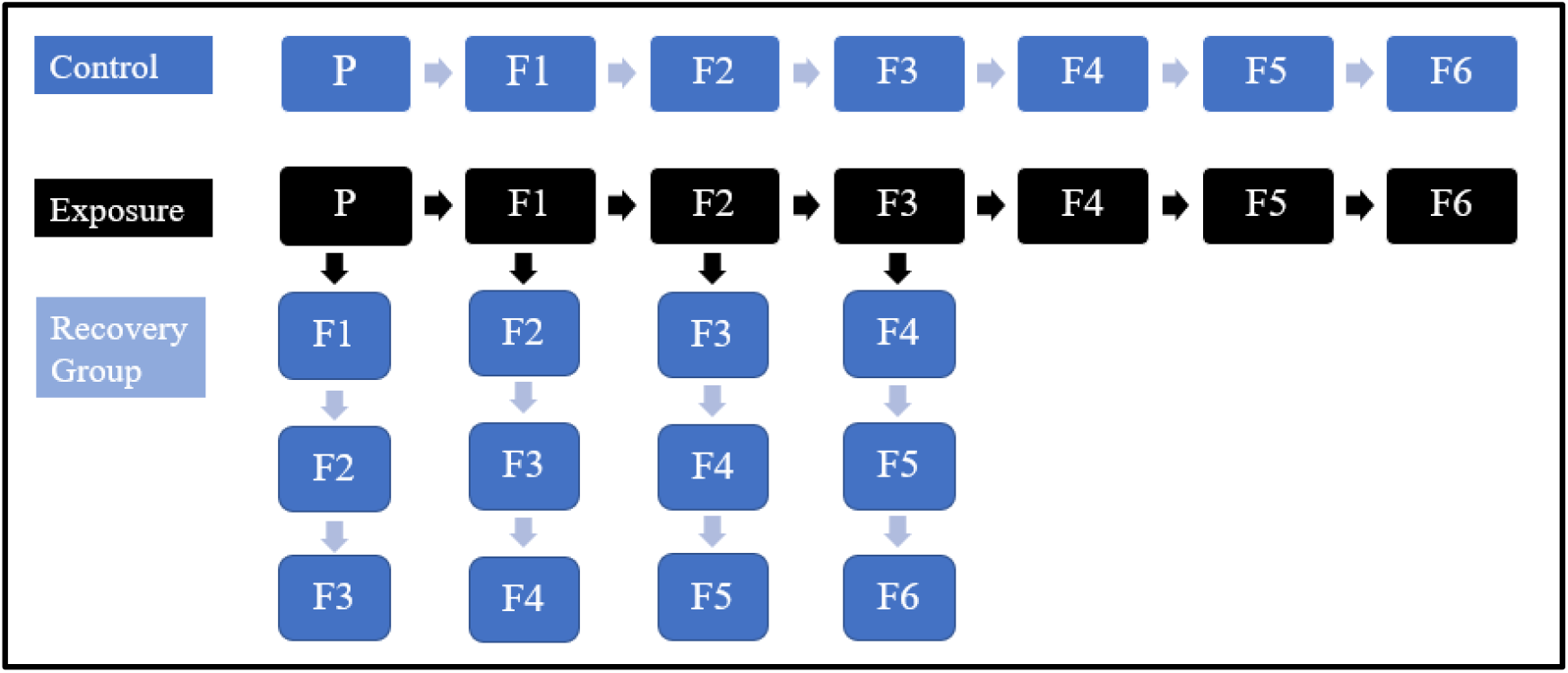
Sketch of intergenerational experiment design. P means parental generation. Blue and black square block represents TWP-free culture medium and TWP leachate medium, respectively.

## 2. Materials and Methods

### 2.1 Preparation of TWP leachate and B. calyciflorus

TWP were purchased from a factory that recycles and smashes scrap tires into particles as artificial turf material. These TWP were sieved through 100 μm nylon net, then sealed and stored at 4°C. TWP were soaked with hard synthetic freshwater (96 mg NaHCO_3_, 60 mg CaSO_4_, 60 mg MgSO_4_ and 4 mg KCl in 1 L distilled water) (USEPA, 1985) at 2500 mg/L in glass bottle and mixed uniformly. The mixture was aerated continuously for 15 days in dark condition at 25°C. It has been demonstrated that there is equilibrium between water and TWP for most components after 14-days leaching (Capolupo et al., 2020; Selbes et al., 2015). TWP leachate were obtained by sieving the mixture through 38 μm nylon net, and frozen at -20°C for temporary storage.

Rotifer *B. calyciflorus* clone strain was established from resting egg that was presented by Professor T.W. Snell of Georgia Institute of Technology, USA. The above hard synthetic freshwater was used as culture medium. The culture condition was kept constantly in illumination incubator with 16L:8D light schedule with light intensity of 4000 lux at 25°C. Rotifers were fed with 5×10^6^ cell/mL *Chlorella pyrenoidesa*, and the medium was replaced every day. Before experiment, some rotifer with amictic egg were chosen from clone strain and synchronized (Kaneko et al., 2016).

### 2.2 Concentration effect of TWP leachate on single generation

After synchronization of parent, the individuals with amictic eggs were pick out. Then, neonate born within 4 hours were put into 24-well plate. There was one individual in one well with 1 mL culture medium. Culture medium with different TWP leachate concentrations (0, 250, 500, 750, 1000, 1500 mg/L) were obtained by diluting the leachate stock of 2500 mg/L with hard synthetic freshwater. The food density was kept at 5×10^6^ *C. pyrenoidesa* cell/mL. Culture condition was the same as that of their parent. Each treatment included 48 individuals. Survival and offspring number were checked every 12 hours, neonates were taken out until all individuals died.

### 2.3 Multi and trans-generational toxicity of TWP leachate on reproduction

Based on the result of above concentration effect on a single generation, two concentrations were chosen to further explore effects of TWP leachate on reproduction during long term multi and trans-generational exposure. The lower concentration did not impose adverse effect on reproduction of first generation, and the higher one exhibited inhibition.

This experiment consisted of three groups, including control, exposure, recovery group (transfer to no-exposure medium) (Fig. 1). After synchronization, neonates born within 4 hours were put into 24-well plate as parental generation. There was one individual in one well with 1 mL culture medium. Other culture conditions keep same with above part. Offspring were picked up as F1 every 12h and transferred into another plate with new culture medium, and so forth until F6. Offspring in exposure treatment were separated into two parts, one of which was continuously exposed as exposure group to verify multi-generational effect, another of which was put into leachate-free medium as recovery group. And recovery groups were maintained further two generations to explore trans-generational effect. In recovery group, the format of treatments name was defined as m-Ex + n-CK, and m and n represented the number of generations. That means that generation-passage line they belong to have suffered m-generation(s) TWP exposure (Ex) then n-generation(s) TWP-free medium (CK). To eliminate difference of brood batch of offspring, the establishment of each generation was divided into 4 times (first 4 broods), 12 individuals from each brood. Each generation of every treatment included 48 individuals. Survival and offspring number was checked every 12 hours until all individuals died.

### 2.4 Statistics

The average individual reproduction and lifespan of rotifers at treatment were calculated by dividing the total offspring number and survival time by individual numbers (picking out missing individual). To display effects of treatment more directly, relative value to control group (%) of average individual reproduction at different treatments was utilized within same generation. Positive value means promotion effect, otherwise inhibition.

## 3. Results

The direct exposure to TWP leachate affected lifespan and reproduction of single generation with dosage-dependent manner, which showed hormesis at low dosage and inhibition at high one (Fig. 2). The 250 and 500 mg/L were chosen as low and high dosage in following multigenerational research. And the reproduction was more sensitive to TWP leachate than lifespan, thus reproduction was used only as evaluation index in following parts.

**Fig. 2.**
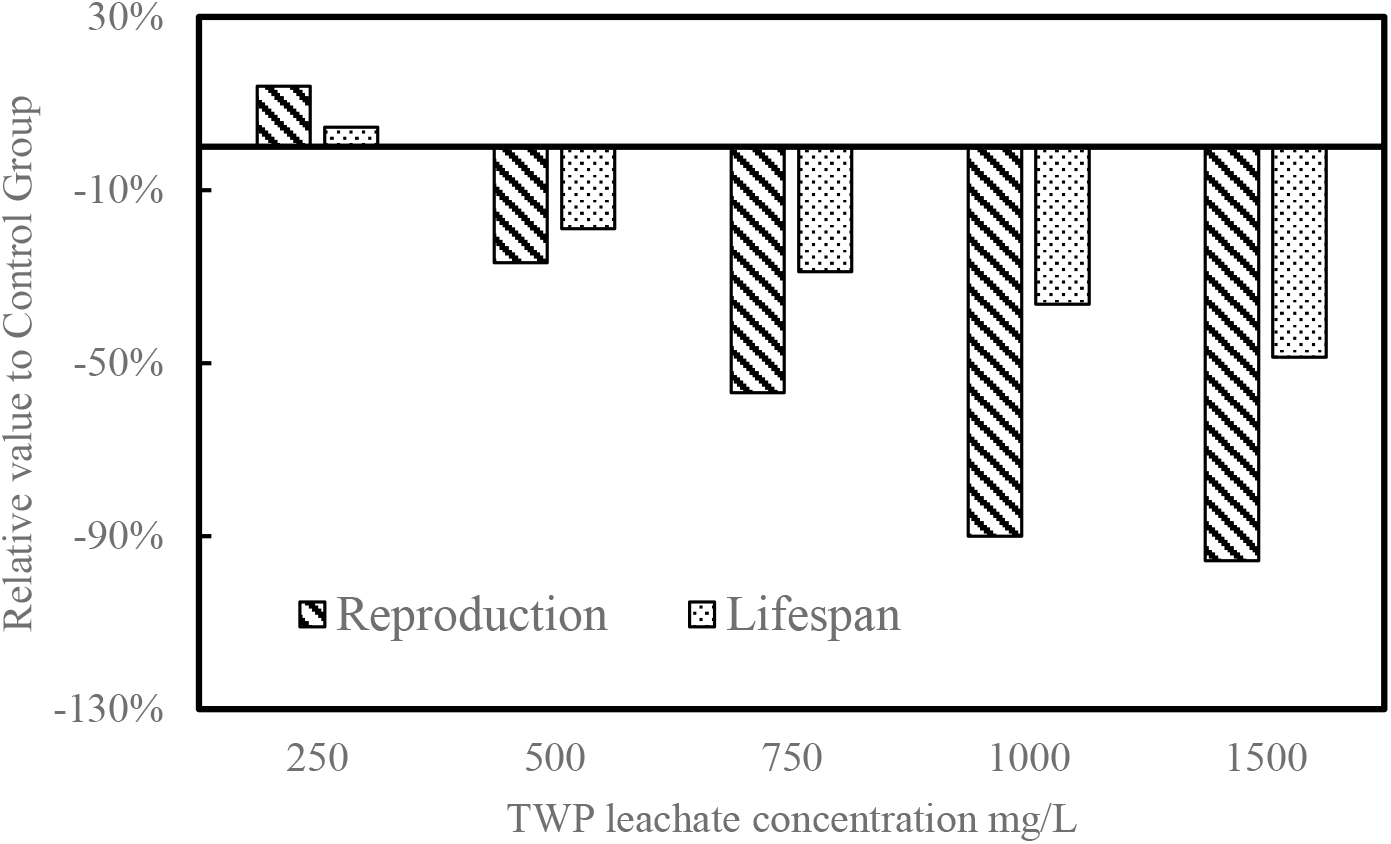
Relative value to control group for average individual reproduction and lifespan at different TWP leachate concentrations. Background slash and dot represents reproduction and lifespan, respectively.

The persistent exposure to TWP leachate gradually reduced relative number of offspring per female across generations (Fig. 3), indicating carryover effects of past exposure lowering the resistance to TWP leachate. At low dosage, the transformation of effect from hormesis to inhibition occurred after F2. The population became extinct after F4 by 100% reproduction inhabitation at high dosage. These implied the intergenerational accumulation of carryover effects from past-generations exposure of TWP leachate.

**Fig. 3.**
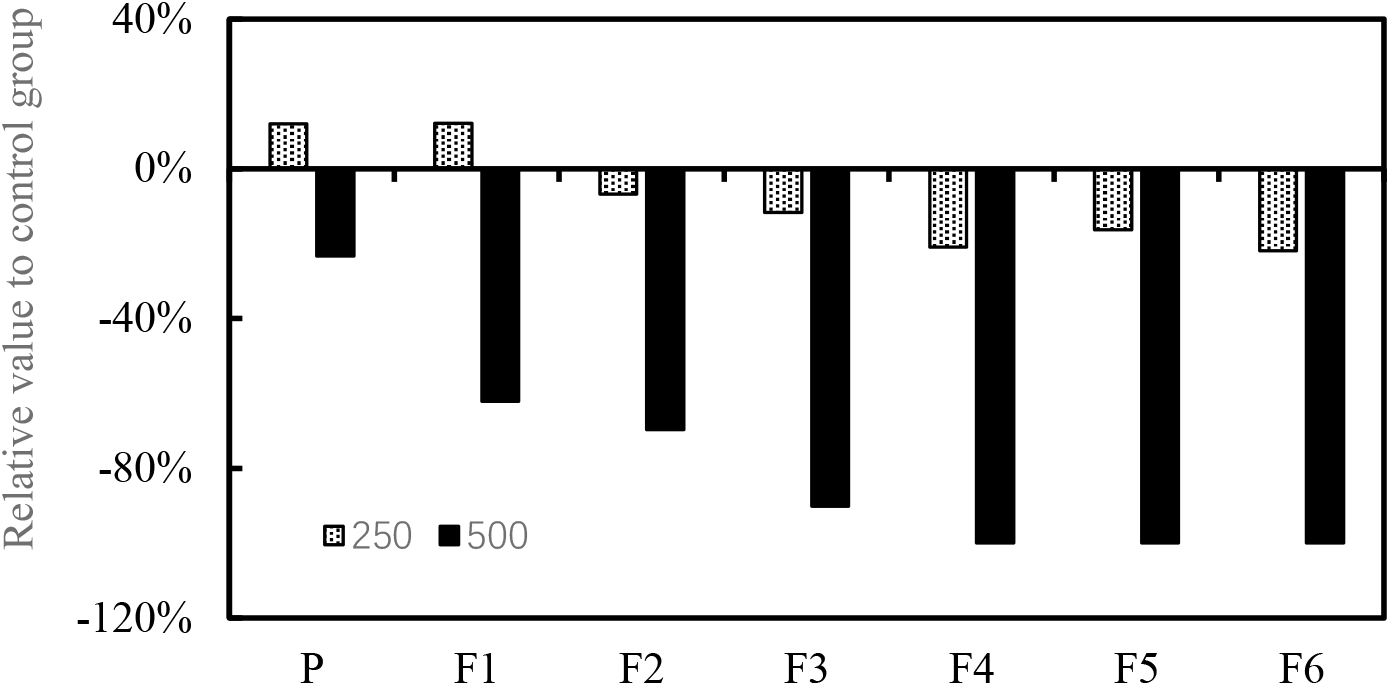
Relative value to control group for average individual reproduction under 250 and 500 mg/L TWP addition at different generations (P – F6). Background dot and black represents 250 mg/L and 500 mg/L, respectively.

When the offspring born from exposure treatment were cultured for 3 continuous generations at no-exposure medium as recovery line, their fecundity trends related to Control were similar with their ancestors exposed to TWP leachate (Fig. 4). It indicated that the effects from parental exposure to TWP leachate could be carried over to their offspring without exposure.

**Fig. 4.**
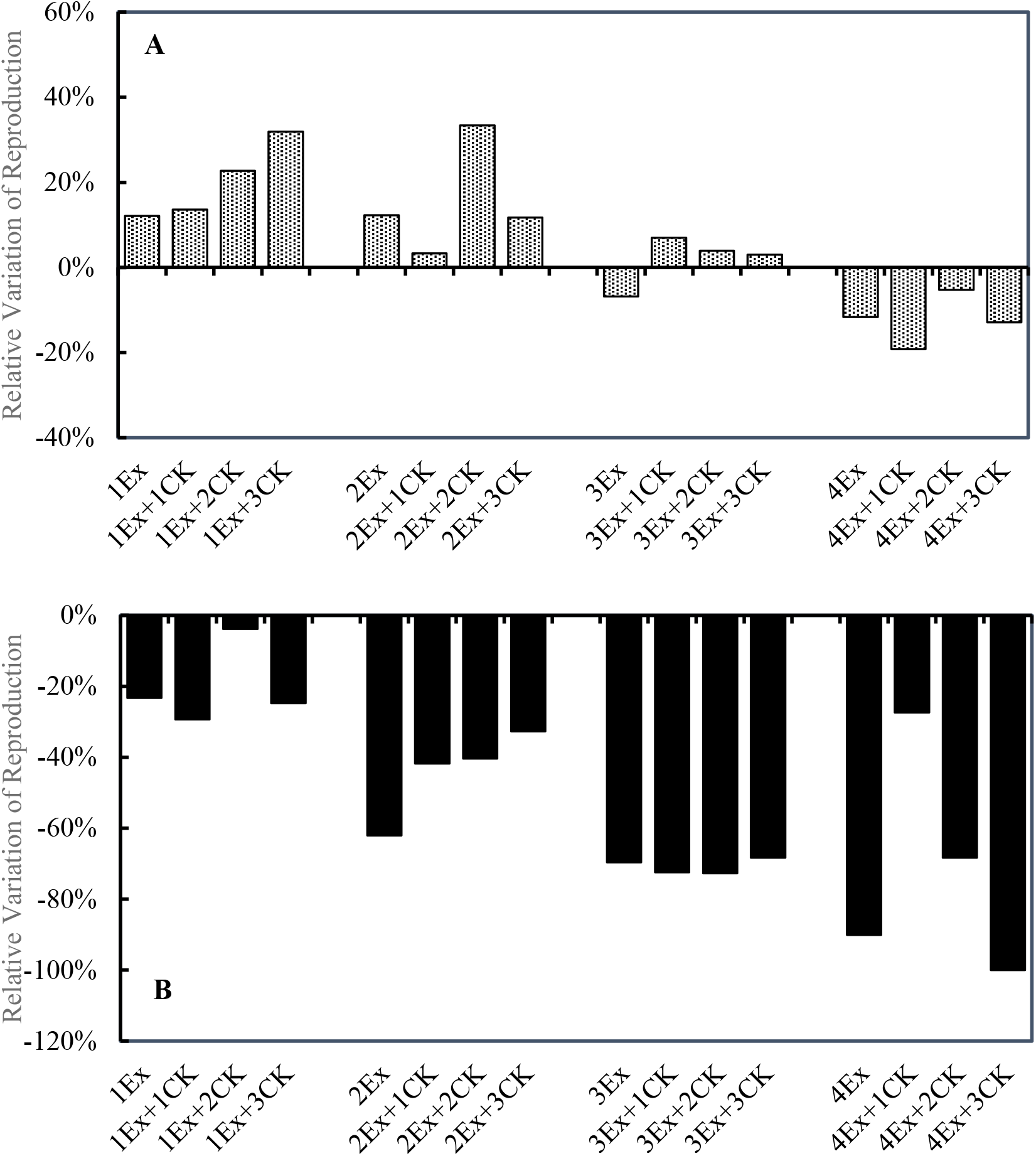
Variation relative to control group for average individual reproduction at different generations transferred neonate into no-exposed medium. A. Transferred from 250 mg/L TWP leachate treatment. **B**. Transferred from 500 mg/L TWP leachate treatment. The format of treatment name, (m) Ex + (n) CK, means that generation-passage line they belong to has suffered m-generation(s) TWP exposure then n-generation(s) TWP-free medium.

## 4. Discussion

Rotifer, as a widespread zooplankton in aquatic ecosystem, build population by quick reproduction and generation passaging. The sporadic road runoff could often expose rotifer to TWP leachate at their specific stage of life cycle and generations. Although it has been shown that TWP leachate exposure influence some zooplankton on a single generation, the transitivity of effects from one generation to their offspring with exposure (or not) are still unknown, principally including additive impacts, adaptability or recovery. We explored these delayed effects based on reproduction performance along generations lineage, built quickly through parthenogenesis. Rotifers exposed to TWP leachate, both to high-dosage and multigenerational low-dosage, generated fewer offspring compared to these without exposure. The TWP leachate exposure from past generations had an additional and extended negative effects on reproduction performance. The carryover effect may tip population into decline even premature collapse over lasting implication.

The persistent exposure of TWP leachate aggregated the negative impact on reproduction. Under low dosage, the effect transferred from hormesis into inhabitation after F2. For high dosage, inhibition rate showed increasing trend along generations, and reproduction halted totally after F4, which predicts that population would develop towards collapse finally. On the other hand, past exposure reduced the current resistance to this stressor. This indicates that past exposure to previous generations could be carried over to offspring by additive manner. This additive effect along generations also has been observed for some other contaminants, which could be attributed to the accumulation *in vivo* and intergenerational transmission of toxic substances.

The cumulative expression of effects observed in this study could give impetus to re-examining safe concentration, which have shown no negative effect on single generation in other zooplankton. Our finding emphasizes the necessity of cautiously interpreting the lack of effects based on original generation in consideration of this carryover effects of stressors. Studies where no effects of TWP leachate exposure for single generation are found may have neglected negative effects that may appear gradually in offspring.

When offspring generated from exposed individual were transferred into culture medium without TWP leachate for continuous 3 generations, regarded as meaningful no-exposed generation in recovery line, the third generation still showed the similar response reproduction with their “great-grandmother” exposed. This means that the effects of TWP leachate can be carried over to offspring except through intergenerational residue and transmission. It could be explained by epigenetic modifications, which triggers heritable effects without change of gene sequence. This inertia of toxicity along generations passaging will extend effect of temporary TWP leachate exposure on reproduction from a longer time scale. Subsequently, the population development and stability may be influenced.

Our study reveals that previous exposure to stressors such as TWP leachate has evident carryover effects on individual reproduction performance of offspring. Some studies have investigated effects of TWP leachate in single generation, but there is no one to explore exposure effects across multiple generations in long term. The rotifer, as well as many other zooplankton, build stable population by quick reproduction and generations passaging based on minor original pioneers. In consideration of sporadic invasion characteristic of TWP leachate in environment which may happen at specific one or several generations during population establishment, it is necessary to assess the transmission of effects across generations. In our study, past exposure of TWP leachate, on the one hand, magnified the inhibition on reproduction by additive manner in persistent multigenerational exposure; on the other hand, extended this reproductive toxicity in offspring without exposure. Those carryover effects on individual reproduction may in further disturb population persistence in long term. These results provide new insight to more comprehensive risk assessment of TWP leachate and emphasize the importance of control of road runoff contained TWP. It merits the further studies to analyze of biochemical and molecular mechanism of carryover effect and strategies to alleviate exposure risk of TWP leachate.

## 5. Conclusions

The TWP leachate can affect lifespan and reproduction of rotifer *B. calyciflorus* with obvious concentration effect. Although low dose of TWP leachate improved reproduction of parental and F1 generations as hormetic effect, inhibition emerged from F3 generation due to multi-generational accumulation of toxicity. Meanwhile, this adverse effect was irreversible at F4 of continuous exposure even though it was transferred into TWP-free medium for recovery. This presents evident trans-generaitonal effect of TWP leachate. For high dose, there were more evident multi and trans-generational toxicity throughout all generations, and rotifer population became extinct after F4. It will be more accurate to draw toxic threshold of TWP based on multi-generations rather than a single generation. And the ecological risks of environmental TWP leachate should be assessed in longer time scale. In addition, analysis for evaluation of effects of different components in TWP should be further investigated.

## Acknowledgements

This work was supported by the Graduate Research and Innovation Project of Jiangsu Province (KYCX20_1190).

## Author contributions

**Yanchao Chai:** Conceptualization, Methodology, Writing - original draft, Funding acquisition. **Haiqing Wang:** Conceptualization, Writing - review & editing. **Mengru Lv:** Investigation. **Jiaxin Yang:** Supervision, Writing - review & editing.

## Declaration of competing interests

The authors declare that they have no known competing financial interests or personal relationships that could have appeared to influence the work reported in this paper.

## Notes

### Competing Interest Statement

The authors have declared no competing interest.

